# Programmable cell differentiation in budding yeast uncouples reproductive and metabolic tasks

**DOI:** 10.64898/2025.12.04.692473

**Authors:** Jacob Guidry, Grant R Bowman

## Abstract

High-yield biomanufacturing requires large cell populations and a mechanism for directing metabolic resources towards product synthesis. However, the resources that support population growth are the same as those that drive productivity, creating a conflict that limits production yields. To overcome this fundamental limitation, we apply the principle of division of labor to separate reproductive and metabolic tasks into distinct cell types within an isogenic *Saccharomyces cerevisiae* culture. We introduce MiSTY (Microbial Stem Cell Technology in budding Yeast), a genetic platform that exploits natural asymmetric cues to control cell differentiation. Leveraging bud cell-specific transcription, a sequential series of recombinase-based genetic circuits generates Activated Stem Cells (ASCs) that divide asymmetrically into two cell types: bud cells that terminally differentiate into Factory Cells (FCs) and mother cells that remain self-renewing ASCs. Time-lapse microscopy demonstrated 100% differentiation fidelity across 97 cell divisions. Phenotypic and genotypic analyses showed that stem cell populations could be converted to over 95% FCs within 24 generations. By converting FCs into leucine auxotrophs, we inhibited FC proliferation while allowing continued ASC division, demonstrating complete uncoupling of cell growth from product synthesis. Because they continuously generate healthy new FCs, MiSTY cultures maintain high levels of productivity even under conditions that severely impair the growth and biosynthetic capacity of metabolically exhausted factory cells.

## Introduction

Industrial biomanufacturing pushes cells beyond their evolved resource capacities, creating a fundamental conflict between growth and productivity that ultimately limits product yields and reduces the potential impact of these technologies (Wu *et al*, 2016),(Mao *et al*, 2024),(Dinh & Maranas, 2023). Cellular maintenance, DNA replication, stress responses, and metabolite synthesis all draw from the same finite pools of ATP, biosynthetic intermediates, ribosomes, and transcriptional capacity (Weiße *et al*, 2015),(Sabi & Tuller, 2019). Industrial biomanufacturing intensifies these demands, imposing levels of metabolic burden that conflict directly with evolved resource allocation priorities (Cordell *et al*, 2023), (Zieringer *et al*, 2021). Devoting cell resources to product formation inhibits biomass accumulation, while investing resources in growth reduces biosynthetic throughput. This fundamental conflict restricts product titers and limits the broader technological impact of microbial biomanufacturing.

In addition to reduced metabolic efficiency, sustained production imposes evolutionary pressures that erode engineered function over time. Accumulation of pathway intermediates or toxic products can impair cell viability, and chronic production stress drives elevated mutation rates that select for variants that down-regulate, disrupt, or entirely lose engineered expression (Arbel-Groissman *et al*, 2023). Long-term continuous culture studies have repeatedly shown that non-producing “cheater” mutants often out-compete producers, since growth advantage is immediately rewarded in well-mixed microbial populations (Rugbjerg *et al*, 2018). This instability is particularly severe in systems where production burdens every cell equally, whether through T7 RNAP–driven pathways, plasmid-encoded expression cassettes, or constitutive metabolic overexpression (Williams & Murray, 2022),(Glass *et al*, 2024).

Numerous metabolic engineering strategies have attempted to address this constraint (Chai *et al*, 2025). Pathway balancing (Àvila-Cabré *et al*, 2025), host metabolism optimization (Zhu *et al*, 2024),(Wang *et al*, 2017), and precursor supply enhancement (Park *et al*, 2023) have improved yields for specific compounds, while inducible systems (Park *et al*, 2023) and dynamic control schemes can temporally separate biomass accumulation from production (von Stosch *et al*, 2016), (Deng *et al*, 2025). Although valuable, these approaches typically require extensive re-optimization for each new metabolic target. A broadly applicable solution that resolves, rather than merely mitigates, the growth–production trade-off remains elusive.

A potential strategy for resolving the conflict in resource allocation is to manage the activities of cells through division of labor. In multicellular organisms, self-renewing progenitors continuously generate differentiated descendants that execute specialized functions incompatible with proliferation (Duckworth, 2021). By segregating reproduction and functional specialization across distinct lineages, this developmental architecture alleviates the growth–performance conflict at its source. Applying an analogous principle in microbial populations could similarly decouple biomass accumulation from biosynthetic productivity.

Recent progress in bacterial systems involves terminal differentiation circuits in *E. coli* that are capable of decoupling replication from biosynthesis. By generating short-lived producer cells from stable progenitors, these systems were able to increase production levels and dramatically delay evolutionary takeover by non-producing mutants (Williams & Murray, 2022),(Glass *et al*, 2024).

Mushnikov and colleagues applied this concept by constructing a stem cell–factory cell architecture in *Escherichia coli* (Mushnikov *et al*, 2019). In their system, proliferative “stem” cells continuously generate terminally differentiated “factory” cells dedicated to biosynthesis while shielding the reproductive population from pathway toxicity (Mushnikov *et al*, 2025), enabling an over eight-fold increase in product titer. While these results established division of labor as a viable strategy for biomanufacturing in bacteria, analogous systems for other industrially relevant microbes, including yeasts (Hong & Nielsen, 2012),(Kulagina *et al*, 2021),(Parapouli *et al*, 2020), remain underdeveloped.

In recent years, genetic engineers have developed methods for engineering consortia of yeast strains, where different strains execute complementary biosynthetic tasks while also influencing their partner strains’ growth rates (Peng *et al*, 2024),(Roell *et al*, 2019). In these co-culture studies, partitioning metabolic pathways across multiple auxotrophic partners substantially reduces the biosynthetic burden on individual strains, improving growth, stabilizing community behavior, and enhancing production of high-value compounds such as resveratrol (Aulakh *et al*, 2023),(Heng *et al*, 2024). This work validates metabolic division of labor as a powerful strategy for reducing burden and improving robustness.

However, these advancements in co-culturing technologies also highlight the intrinsic limitations of multi-strain approaches. Community composition is highly sensitive to initial population ratios and molecular exchange rates, often requiring extensive tuning and external modulation to maintain the desired balance of cell types (van Aalst *et al*, 2023), (Ronda & Wang, 2022). As additional strains are added, these systems become increasingly complex (Jiang *et al*, 2023). These constraints underscore a need for division-of-labor strategies that preserve the advantages of metabolic partitioning while avoiding the liabilities of multi-strain community management.

Afonso, Silver, and Ajo-Franklin demonstrated that native asymmetry in *Saccharomyces cerevisiae* could be exploited to direct cell differentiation within an isogenic cell population (Afonso *et al*, 2010). They developed an Ace2-dependent pathway for driving expression of a cytotoxic enzyme exclusively in bud cells, and showed that its inhibitory effects could be confined to only one of the two daughter cell progeny. This work is foundational to the method for decoupling the tasks of cell growth and product synthesis that we describe.

Here, we present MiSTY (Microbial Stem Cell Technology in budding Yeast), a platform that implements stem cell–supported division of labor in *S. cerevisiae* by combining native asymmetric division machinery with synthetic lineage-control circuits. MiSTY establishes three functional cell states: Inactive Stem Cells (ISCs) that proliferate without differentiating, Activated Stem Cells (ASCs) that undergo asymmetric budding to continuously renew themselves while producing specialized progeny, and Factory Cells (FCs) that terminally differentiate to perform biosynthesis. Cell fate transitions are executed through sequential, irreversible recombinase-mediated genomic rearrangements, generating stable lineage relationships in which ASCs act as durable long-term progenitors that support sustained FC production.

## Results

### Platform Design and Genetic Architecture

In conventional biomanufacturing approaches, a uniform population of “factory cells” are instructed to commit themselves to product synthesis (Xin *et al*, 2019),(Lindskog, 2018). However, highly productive factory cells often experience metabolic burdens that lead to cytotoxicity, and this limits the number of factory cells that can be generated and the final product yield (Wu *et al*, 2016) (Figure 1A). This limitation may be overcome by dividing the tasks of cell division and product synthesis between two differentiated cell types (Williams & Murray, 2022),(Aulakh *et al*, 2023). In *S. cerevisiae*, this could be accomplished by activating a genetic program that differentiates mother cells from bud cells, such that mothers retain the capacity to divide while their progeny differentiate into growth-limited factory cells (Figure 1B).

**Figure 1.**
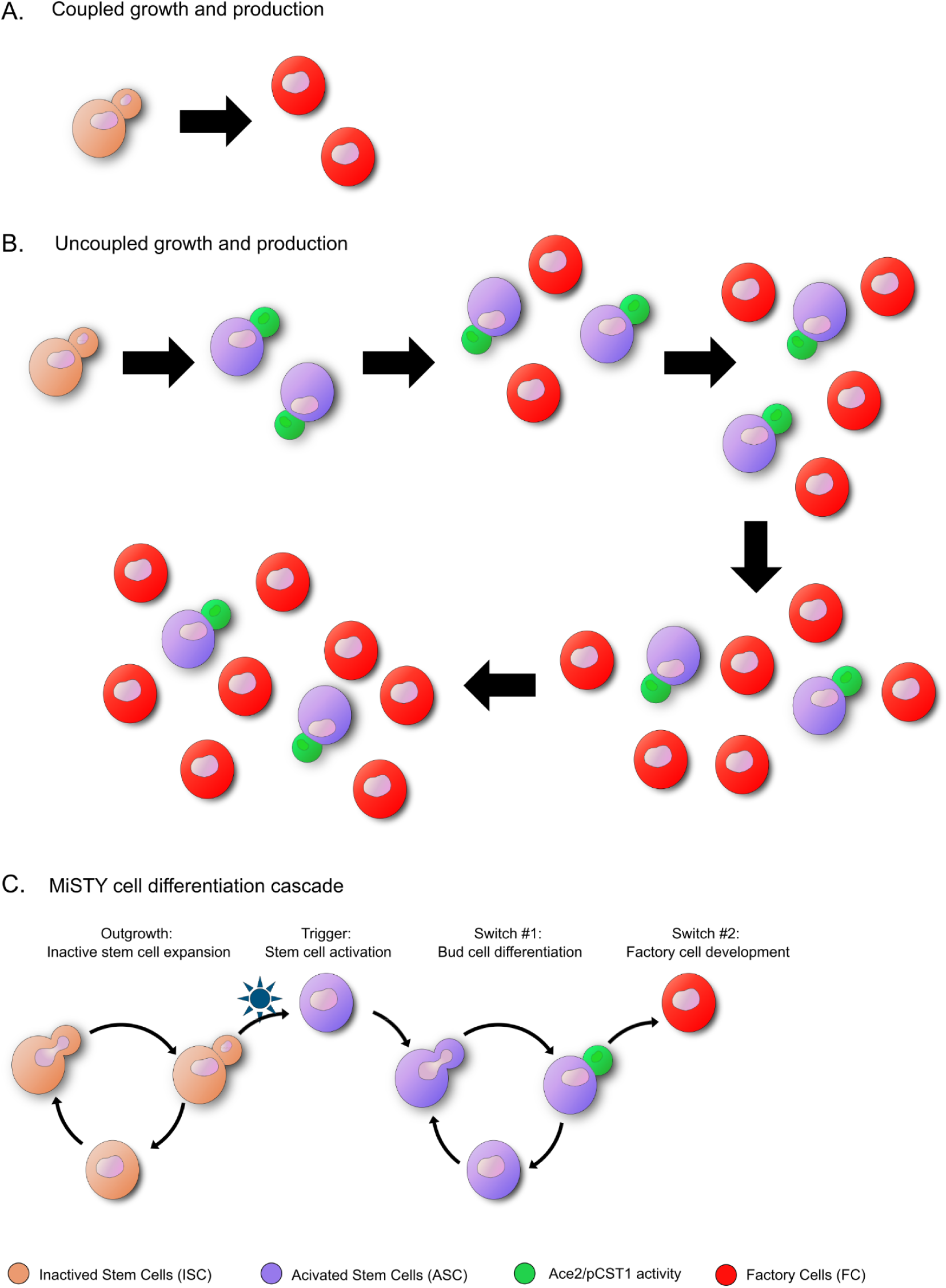
Microbial Stem Cell Technology in Yeast (MiSTY) A) An example of conventional product synthesis, where biosynthetic activity inhibits cell growth. B) Stem Cell-supported synthesis, where stem cells divide asymmetrically into one biosynthetically active cell and one stem cell. Even if production inhibits cell growth, the cell population increases in conjunction with product synthesis. C) The MiSTY program for programmable division of labor, where an inducer triggers an asymmetric cell division that leads to cell differentiation and the uncoupling of reproductive and biosynthetic tasks.

Our strategy for engineering inducible asymmetric cell division and cell differentiation in *S. cerevisiae* was to adapt natural mechanisms for differentiating mother cell and bud cell gene expression programs (Dohrmann *et al*, 1996). The goal was to make the cell differentiation event inducible (Trigger, Figure 1C) and to redirect the natural pattern of cell asymmetry toward programmed cell differentiation. A core functional component of this engineered system is the Ace2 transcription factor, which achieves asymmetry through bud cell-specific nuclear localization (Voth *et al*, 2007),(Herrero *et al*, 2020). Following the work of Afonso et al. (Afonso *et al*, 2010), we connected the activity of Ace2 to a cell fate decision by engineering the output of the *CTS1* promoter (p*CTS1*), which contains multiple Ace2 binding sites and is therefore active exclusively in bud cells during late mitosis and early G1 phase. This expression window (Switch #1, Figure 1C) aligns precisely with our design requirement for asymmetric fate specification. By placing a site-specific recombinase gene under p*CTS1* control, we could achieve bud cell-specific genomic rearrangements to drive irreversible commitment to Factory Cell fate (Switch #2, Figure 1C).

### A Bud Cell-Specific Recombinase

We employed Cre as the site-specific recombinase under control of p*CTS1*. Mindful that Cre must be targeted to the nucleus in order to function and that excessive or prolonged Cre activity could have deleterious off-target effects (Sauer, 1993), we added regulatory elements to Cre that drove its nuclear localization and timely degradation as the bud cell entered G1/S transition. Here again, we followed the work of Afonso et al. (2010), who used these elements to control the timing and localization of a cytotoxic enzyme. The combination of a Nuclear Import and Nuclear Export Signal (NLS and NES) drives shuttling between cytoplasm and nucleus (Wen *et al*, 1995), and a Sic1 degradation tag promotes rapid degradation via the 26S proteasome after the protein is phosphorylated and subsequently ubiquitinated at the G1/S transition (Petroski & Deshaies, 2003),(Cross *et al*, 2007). To visualize the transient expression, nuclear localization, and subsequent degradation of Cre, we also included a GFP tag. The final fusion architecture of this construct, named p*CTS1*-Cre, was Sic1deg-NES-GFP-Cre-NLS (Figure 2A).

**Figure 2.**
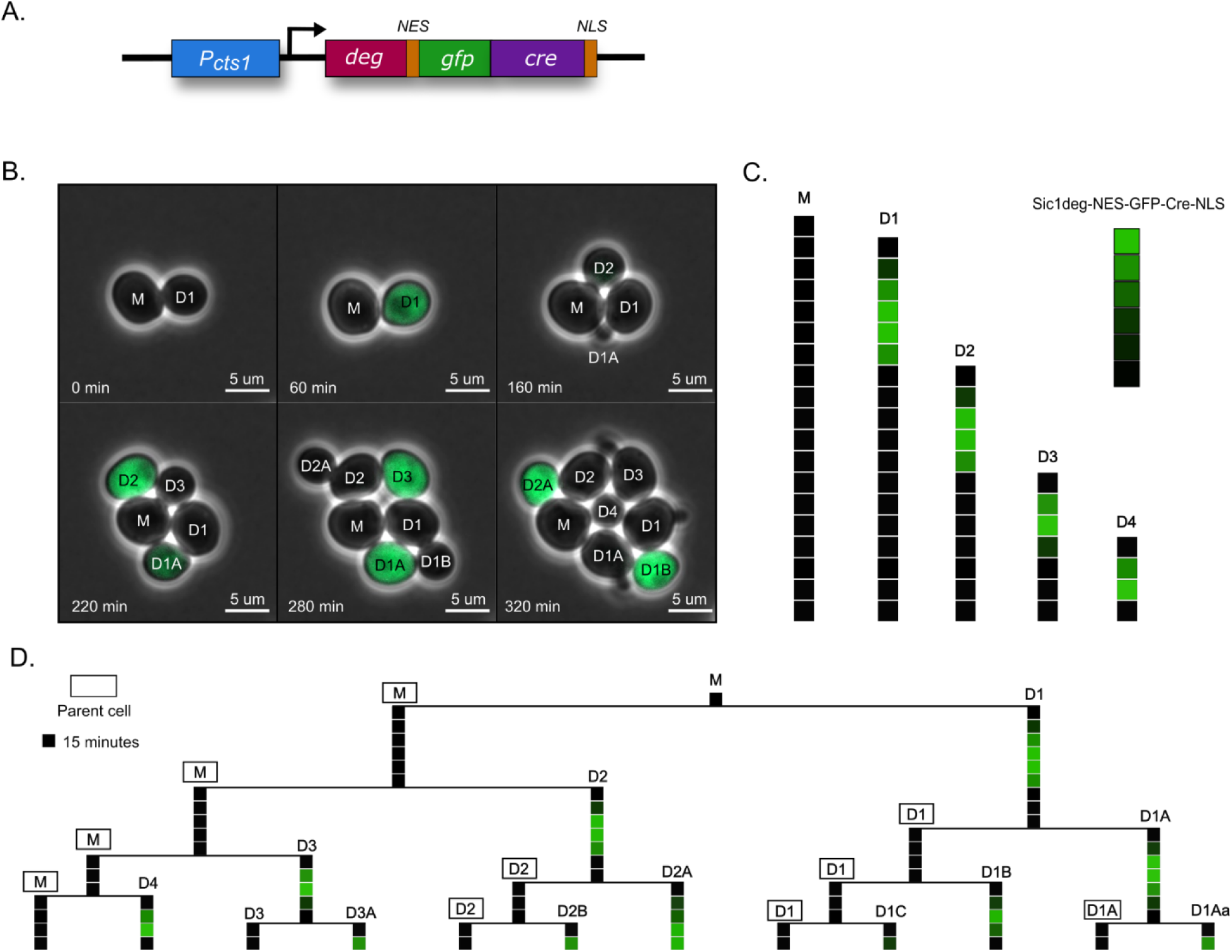
Temporal and Spatial Control of Cre Recombinase. A) The engineered p*CTS1*-Cre gene for driving bud cell differentiation. B) Time-lapse imaging of AA311, observing the cell cycle-dependent localization of the Cre fusion protein. M marks the mother cell and D marks bud cell daughters, with numbers 1-4 indicating the generation born from the mother cell and the secondary letters indicating the order of grandchildren born from the daughters. C) Representations of the average GFP fluorescence intensities of selected cells in B, showing one transient period of GFP fluorescence in the history of each cell. D) Representations of the average GFP fluorescence intensities of all cells in B, with lineages organized as a tree and cell division events represented by horizontal bars. For C and D, each square panel represents a snapshot after one 15 minute interval, with time increasing from top to bottom.

After integrating p*CTS1*-Cre into a normal yeast genetic background (Strain AA311), we followed the localization pattern of the Cre fusion protein in individual cells using time-lapse fluorescence microscopy (Supplementary Video V1 and Figure 2B). GFP-Cre signal appeared exclusively in bud cells, during late stages of bud development or soon after septation. This signal peaked approximately 15-30 minutes after cytokinesis, then diminished over the following 30-90 minutes, consistent with active degradation during G1/S (Figure 2C). The pulse of GFP-Cre fusion protein occurred only once in each cell’s lifetime and never recurred, even as those cells matured and produced their own daughters (Figure 2D). This pattern demonstrates that the p*CTS1*-Cre gene product exhibits bud cell specificity and transient one-time accumulation.

To assess the Cre fusion protein’s ability to specify Factory Cell fate determination in budding cells, we constructed a recombination-based reporter gene for Cre recombinase activity. Here, the constitutive p*TEF1* promoter is placed upstream of the red fluorescent reporter mChy, but mChy expression is blocked by the insertion of a *LEU2* coding sequence and a strong transcriptional terminator downstream of the promoter. *LEU2* and the terminator are flanked by *loxP* sites placed in direct orientation, such that nominal Cre recombinase activity deletes these elements and places the constitutive promoter immediately upstream of mChy (Switch #2, Figure 3A). When we genomically integrated the reporter gene into a normal yeast genetic background (Strain AA320), the strain was not fluorescent, as expected (Supplementary Figure 1A) Fig. When we added p*CTS1*-Cre (Strain AA321), all cells were bright red, indicating that the Cre fusion protein is capable of activating this reporter (Supplementary Figure 1B).

**Figure 3.**
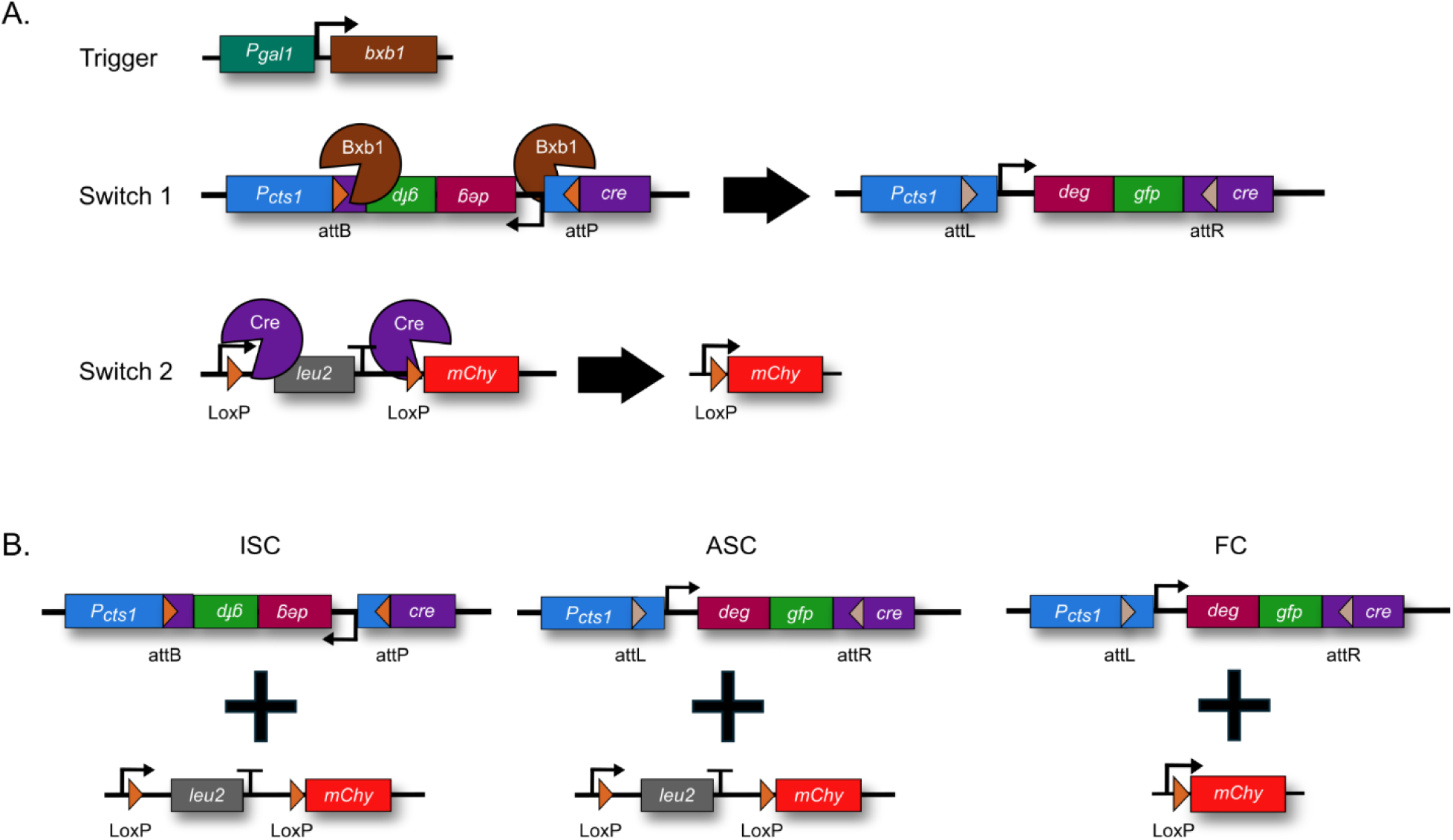
MiSTY Genetic Circuitry. A) Genetic elements in MiSTY. Coding sequences = colored rectangles; promoters = angled arrows; recombinase sites = triangles; and recombinase enzymes = sectored circles. Arrowheads show switches converting from initial to final state. B) Each cell type in the MiSTY developmental pathway is associated with a distinct configuration of Switch #1 and Switch #2.

### A Trigger for Inducing Asymmetric Cell Division

To determine the frequency at which p*CTS1*-Cre activates Switch #2, it was necessary to control the timing of Cre activity. To do this, we engineered an additional genetic switch, wherein Cre expression could be switched from an inactive to an active state. To enable this switch, we employed a second site-specific recombinase, Bxb1, which catalyzes unidirectional recombination between *attP* and *attB* sites, yielding products that are poor substrates for reverse recombination (Olorunniji *et al*, 2017). This irreversibility makes Bxb1 ideal for permanent genetic switches (Sales *et al*, 2025). To create a version of Cre that could be activated by Bxb1 activity, we inverted a central segment of p*CTS1*-Cre and flanked it with *attB* and *attP* sites, creating a construct named Switch #1 (Figure 3A). We placed the site of inversion within a flexible, non-structured region that would likely tolerate the sequence modifications imposed by the recombination site.

To ask if the inverted Cre gene is non-functional, we integrated Switch #1 into a *S. cerevisiae* strain that also contained the Switch #2 reporter (Strain AA715) and observed no GFP or mChy expression, indicating that the Cre fusion protein is not expressed and that there is no Cre recombinase activity. To create a mechanism for activating Switch #1, we placed the Bxb1 coding sequence under control of the galactose-inducible p*GAL1* promoter (Lettow *et al*, 2021), creating a Trigger construct (Figure 3A), which we integrated into *S. cerevisiae* that also contained Switch #1 and Switch #2. The resulting strain (AA639) comprises the complete MiSTY genetic system.

### Observation of Asymmetric Cell Division and Differentiation

According to this circuit design, p*GAL1*-*BXB1* is expected to be repressed when cells are grown in glucose-containing media, and both Switch #1 and Switch #2 should remain in their initial state and the cells should be negative for GFP and mChy fluorescence. We call these types of cells Inactive Stem Cells (ISCs) to reflect their precursor state (Figure 1C and 3B). Upon switching to galactose-containing media, p*GAL1* is expected to drive Bxb1 expression, catalyzing the inversion of Switch #1 and restoring a functional p*CTS1*-Cre expression unit. We call these types of cells Active Stem Cells (ASCs) to reflect their ability to divide asymmetrically and produce progeny with different cell fates. The bud cell progeny of ASCs activate Switch #2, converting them into a terminally differentiated cell type that we call Factory Cells (FCs).

When we observed Strain AA639 after growth in glucose-containing media, we were initially surprised to find that a majority of the cells exhibited bright mChy fluorescence, indicating that they had switched to Factory Cell fate, and suggesting that the differentiation mechanism may be sensitive to leaky Bxb1 expression. To avoid un-induced activation of the genetic circuit, we took advantage of a key design feature in Switch #2, which was the inclusion of the *LEU2* amino acid biosynthesis coding sequence in the DNA that is excised by Cre. After the removal of *LEU2*, all cells in the Factory Cell lineage become leucine auxotrophs, while ASCs and ISCs retain the ability to produce this essential amino acid. When we grew Strain AA639 in glucose-containing synthetic drop-out media lacking leucine (SDglu-Ura), the number of mChy positive cells was reduced to approximately 11% (Figure 5A). This provided a growth condition that would allow us to observe the developmental processes triggered by Bxb1 expression.

**Figure 4.**
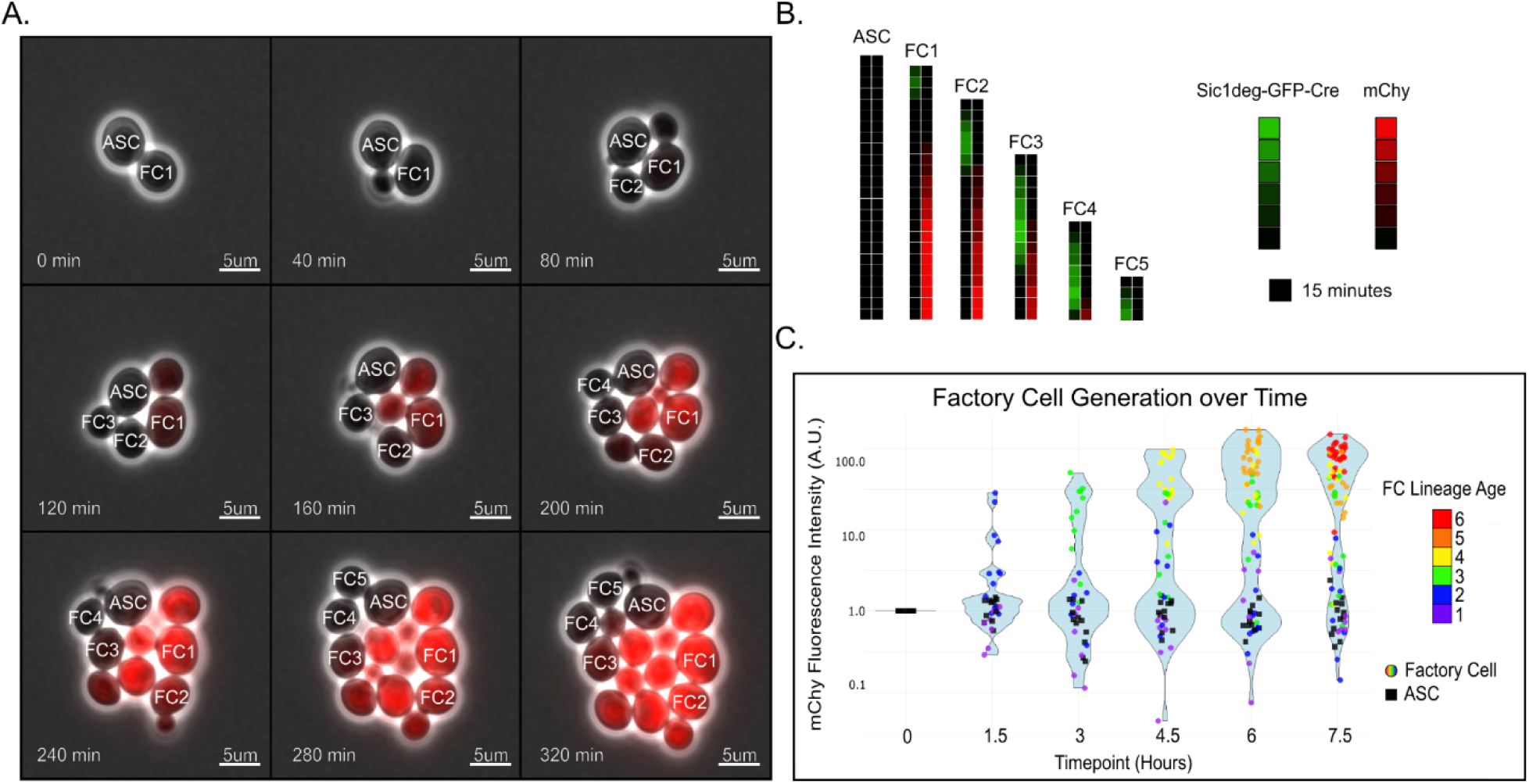
Activated Stem Cell Division and Factory Cell Differentiation. A) Timelapse imaging of AA639, observing the accumulation of mChy fluorescence in differentiated bud cell lineages. ASC marks an Activated Stem Cell and FC marks bud cell progeny, with numbers 1-5 indicating the generation born from the mother cell. B) Representations of the average GFP and mChy fluorescence intensities of selected cells in A, showing one transient period of GFP fluorescence followed by the accumulation of mChy fluorescence. Each square panel represents a snapshot after one 15 minute interval, with time increasing from top to bottom. C) Lineage tracking of 23 different ASCs and their FC progeny over 7.5 hours. ASCs are represented by black circles and circles representing FCs are colored according to the number of mother cell generations away from their birth. Relative mChy fluorescence intensities of individual cells are plotted on the Y-axis. Violin plots (blue shading) show the frequency distribution of data points along the Y-axis.

**Figure 5.**
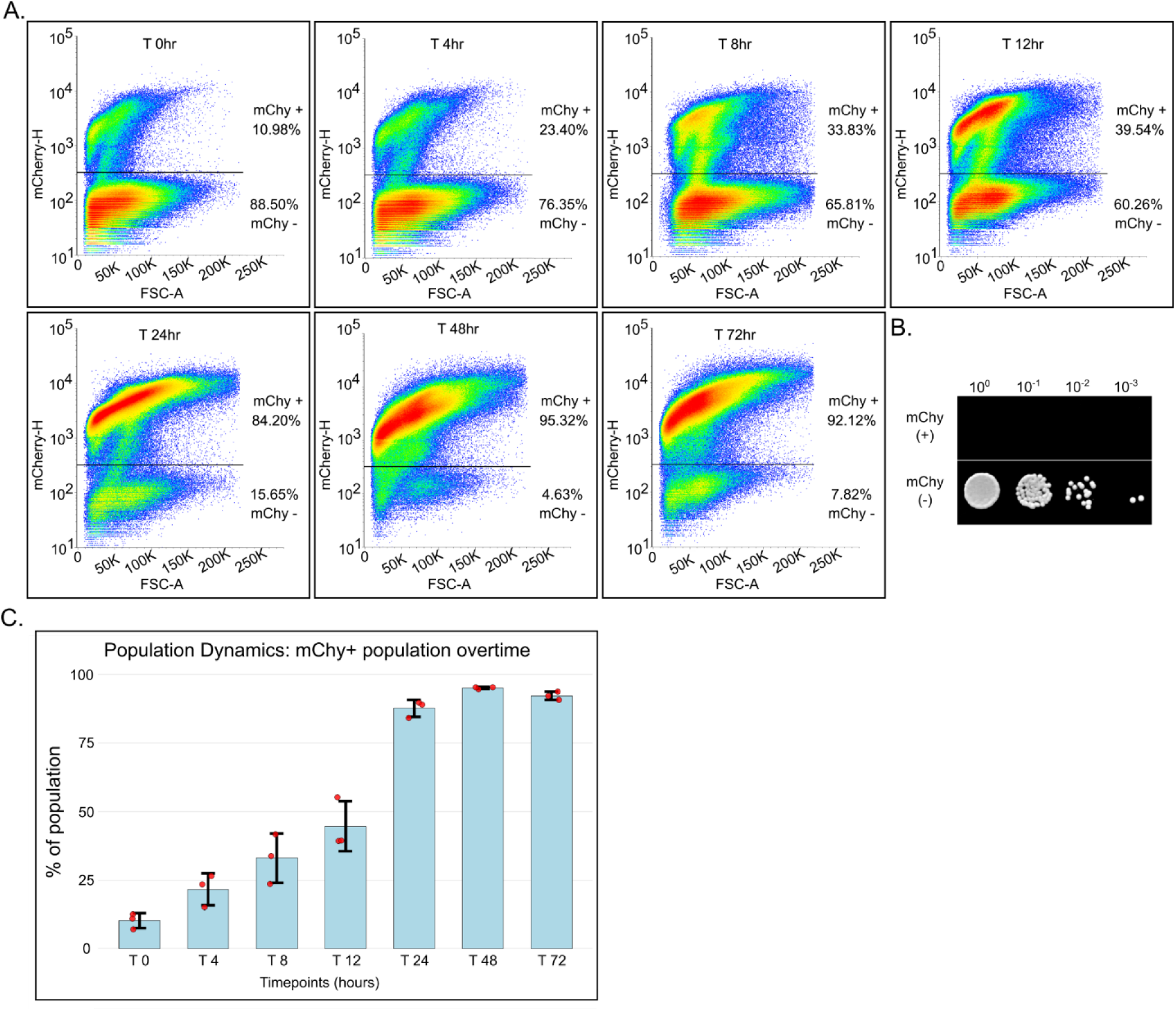
Factory Cell Population Dynamics in Liquid Suspension Cultures. A) Flow cytometry plots depicting mChy fluorescence intensity (Y-axis) as a function of side scatter (X-axis) for individual cells over a 72 hour time course following the induction of immature stem cells from strain AA632. The horizontal line indicates the gate used to distinguish mChy positive from mChy negative cells. B) Equal numbers of mChy positive and mChy negative cells from the 24 hour time point in A were sorted into separate pools, serially diluted, then spot plated onto an agar plate containing leucine-deficient medium. C) Percentages of mChy positive cells over three replicates of the experiment shown A. Bars show standard deviation among three independent trials.

To observe these events, we grew strain AA639 to exponential phase in SDglu-Leu medium, then switched the cells to galactose-containing media lacking leucine (SDgal+glu-Leu) to trigger the conversion of ISCs to ASCs and maintain suppression of the growth of new Factory Cells. Using time-lapse fluorescence microscopy, we observed the growth and development of ASCs over an 8 hour time period (Supplementary Video V2 and Figure 4A-B). Similar to the strain with unregulated p*CTS1*-Cre (Figure 2), the GFP-Cre signal appeared only in bud cells, peaked approximately 15-30 minutes after cytokinesis, and declined in the subsequent 30-90 minute time period during G1/S. In AA639 cultures, the bud cell descendants began to express mChy between 60 and 90 minutes after the pulse of GFP-Cre, indicating that they had differentiated into Factory Cells. The level of mChy signal increased steadily over subsequent hours, consistent with the constitutive expression of the reporter.

We tracked the fates of 23 individual ASCs and their progeny across an average of 4.2 generations per ASC, a total of 97 division events (Figure 4C). Across all divisions, 100% of bud cells showed the expected differentiation pattern: transient GFP followed by sustained mCherry expression. None of the 23 tracked ASCs exhibited GFP or mCherry fluorescence at any time during the observation period. ASCs divided continuously, with doubling times ranging from 120-150 minutes, which matched normal growth rates in galactose-condaining SD medium. Their Factory Cell progeny divided an average of 3 times before arresting, indicating that these newly differentiated leucine auxotrophs are able to grow for some time before the amino acid biosynthesis defect takes effect (Supplemental Figure 2). Overall, we conclude that the processes of asymmetric cell division, Factory Cell differentiation, and ASC maintenance occur at nearly 100% fidelity in galactose-induced MiSTY cultures.

### Population-Level Dynamics

We used multiple phenotypic analyses to follow the changes in cell type composition in liquid suspension cultures. In these experiments, we prepared MiSTY Strain AA639 by growing it to exponential phase in SDglu-Leu medium, then induced the developmental program by switching to SDgal+glu-Leu medium at time zero. We collected samples for analysis at 0, 4, 8, 12, 24, 48, and 72 hours post-induction. To maintain continual growth, samples were diluted into fresh SDgal+glu-Leu to an OD600 of 0.1 every 12 hours.

We measured the mChy fluorescence intensities of individual cells by flow cytometry (Figure 5A), and found that approximately 11% of cells were mChy-positive at time zero, likely reflecting the background rate of Factory Cell differentiation arising from basal Bxb1 Recombinase expression. At t=4 hours, the mChy-positive population increased to 23%, and continued to increase until t=48 hrs, when the percentage of mChy-positive cells surpassed 95%. To ask if fluorescence intensity correlated with Switch #2 recombination, we sorted the mChy positive and mChy negative cells into separate pools and plated them on agar media lacking leucine (Figure 5B). Leucine heterotrophs were abundant in the mChy-negative pool, indicating the presence of ISCs and ASCs. Zero leucine heterotrophs were found in the mChy-positive pool, indicating that all of these cells had passed Switch #2. AA639 cultures behaved consistently across three independent flow cytometry trials (Figure 5C).

To compare the timing of mChy accumulation with the timing of Switch #2, we plated samples from the AA639 induction time course on leucine-supplemented and leucine-deficient agar (Figure 6A-B). In these experiments, the ratio of the number of cells that grow in the absence of leucine compared to the number of cells that grow in the presence of leucine indicates the fraction of cells that have experienced Cre recombinase activity at Switch #2. At time zero, approximately 20% of the cells were leucine auxotrophs, representing the basal level of ISC conversion in galactose-free medium. This is higher than the number of mChy-positive cells in the population, possibly because the agar plating assay counts ASCs and Factory Cells as leucine auxotrophs, while the flow cytometer counts only mature Factory Cells. The percentage of leucine auxotrophs increased rapidly over the t=4 to t=24 hour time period, and ultimately climbed to over 99% of the cell population and remained in this state for the next 3 days. The increase in leucine auxotrophy preceded the increase in mChy fluorescence by roughly 4 hours, consistent with the idea that newly converted ASCs take time to divide and that new Factory Cells require time to accumulate mChy.

**Figure 6.**
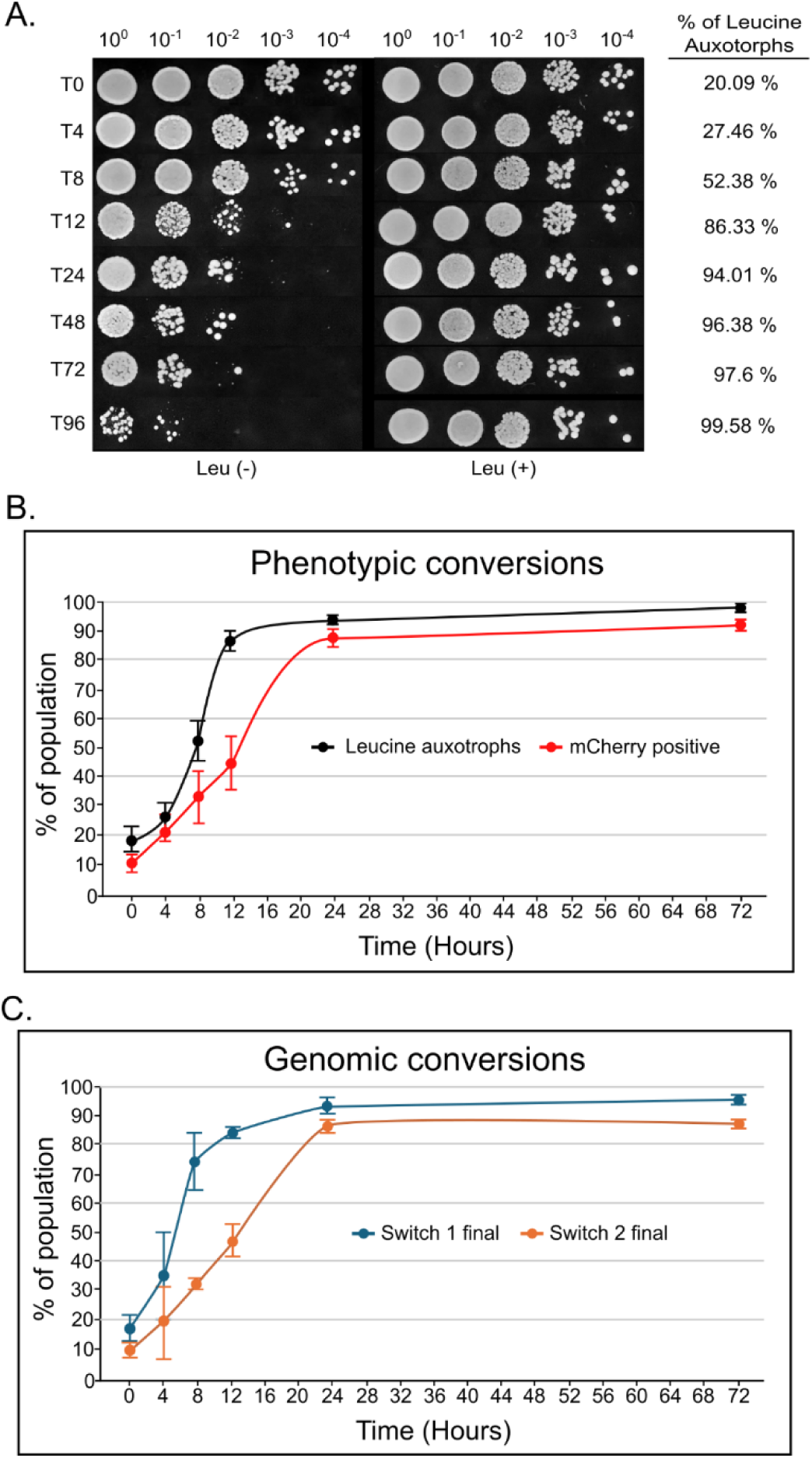
Dynamic Phenotypic and Genotypic Properties of MiSTY Cell Cultures. A) Aliquots of cell culture during a 72 hour time course following the induction of immature stem cells from strain AA632 were serially diluted and spot plated on agar plates, on leucine-deficient (left) or leucine-supplemented (right) media. The percentage of leucine auxotrophs was calculated by dividing the number of colonies growing on Leu- by the number of colonies growing on Leu+ and subtracting from 1. B) Average percentage of leucine auxotrophs over time in A (black line) and mChy positive cells from the same experimental conditions in Figure 5C (red line), with bars showing standard deviation among three independent trials. C) Average percentage of sequence reads indicating the conversion of Switch #1 (blue line) and Switch #2 (orange line) from initial to final state, from cultures collected under the same experimental conditions as in A. Bars show standard deviation among three independently collected datasets.

In our agar plating experiments, we noted the appearance of micro-colonies on leucine-deficient media (Supplemental Figure 3A). Although present at all time points, these micro-colonies became increasingly numerous at t=12h. These cells were unable to sustain growth on leucine-deficient media but grew normally when transferred to leucine-supplemented media (Supplemental Figure 3B). Microscopic examination revealed that all cells expressed mChy, indicating Factory Cell identity (Supplemental Figure 3C). We hypothesize that these colonies originated from ASCs that, while not themselves leucine auxotrophic, gave rise to Factory Cell progeny that retained a limited capacity for growth due to the carry-over of Leu2 enzyme and *LEU2* mRNA that existed at the time of Switch #2 recombination (See also Figure 4A). As microcolonies were not included in our colony counts, these observations support the idea that ASCs are included in the calculated fraction of leucine auxotrophs.

We conclude that the kinetics of leucine auxotrophy align with the onset of mChy fluorescence in a series of events that trace Factory Cell differentiation. We observed near-complete conversion to Factory Cell fate within 24 hours, and sustained levels through 96 hours, confirming long-term stability of the dominant Factory Cell population.

### Genomic Validation of Recombination Events

To measure the timing of the genetic recombination events directly, we analyzed samples from the AA639 induction time course by whole genome sequencing. To obtain reliable quantitative information on the fraction of cells that had undergone recombination events, we analyzed three independent sets of samples from each datapoint, oversampling between 97X and 240X genome coverage. Long-read sequencing enabled unambiguous mapping across recombination junctions, allowing us to distinguish initial and final states at Switch #1 and Switch #2 genetic loci. At time zero, we found that the percentage of sequence reads showing Switch #1 and Switch #2 in their final states was 12% and 8%, respectively (Figure 7), which is consistent with the basal level of circuit activation that we observed in phenotypic assays (Figure 5 and 6). Over the next 24 hours, those percentages climbed to over 90% of total, following the rates of phenotypic conversion. We found little evidence of recombinase errors at the switch points. Cre excision at Switch #2 was always confined to the *loxP* sites, and Bxb1-mediated deletion events at Switch #1 were never observed until the 72 hour time point, when these represented approximately 2% of sequence reads. Notably, the timing of Switch #1 preceded Switch #2 by roughly 4 hours. This is consistent with the difference in timing that we observed in agar plating and flow cytometry assays, and reflects the time that newly generated ASCs need to divide and generate differentiated Factory Cell progeny.

**Figure 7.**
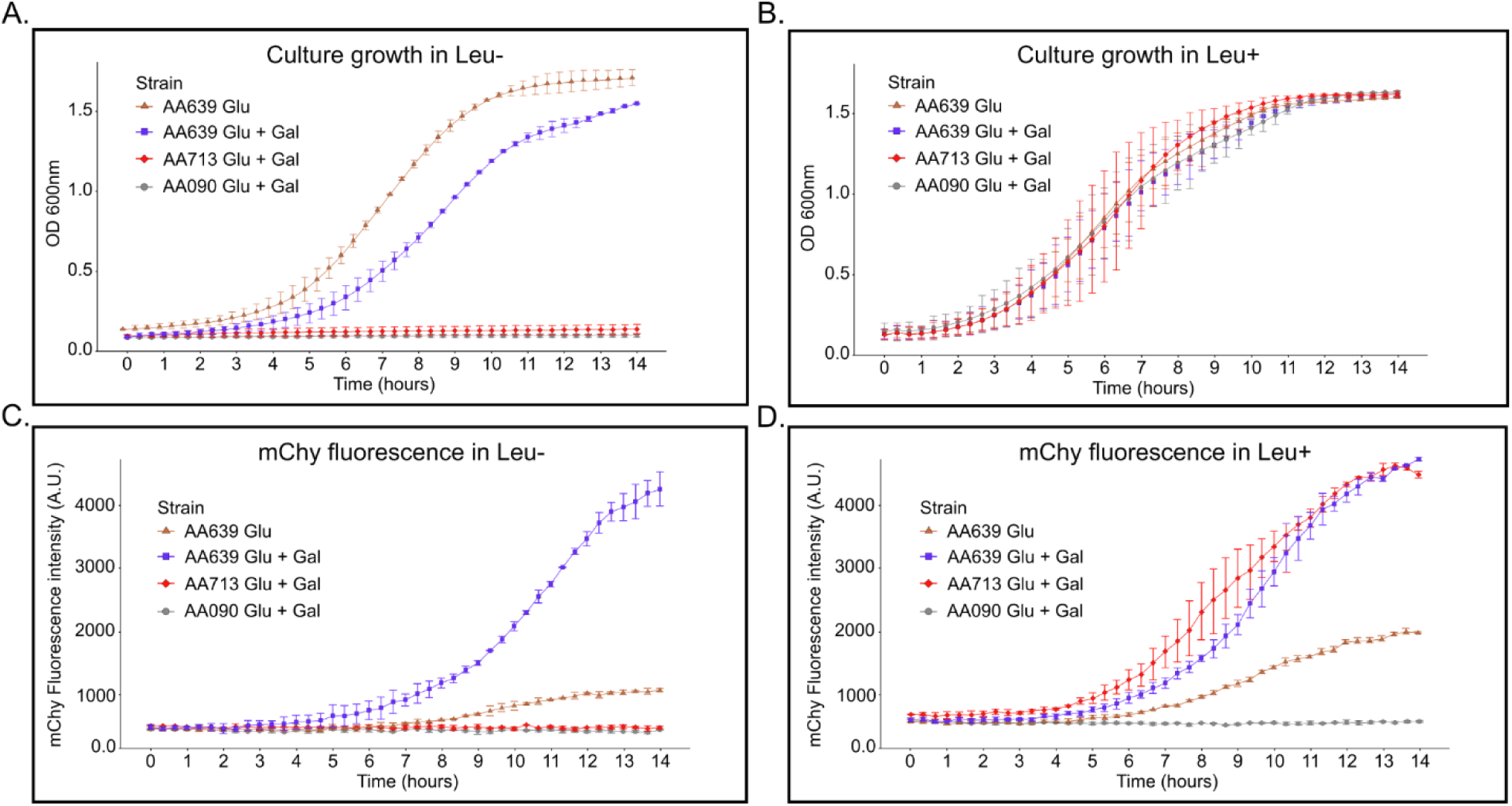
Concurrent Population Growth and Product Synthesis in MiSTY Culturing Conditions. A-D) Exponentially growing strains were cultured in media supplemented with the indicated carbon source for 8 hours, diluted to an OD_600_ of 0.1, then allowed to continue growing in leucine-deficient (left panels) or leucine-supplemented (right panels) media in 96-well plates. Optical density measurements (upper panels) tracked culture growth while Fluorescence detection (lower panels) tracked the increase in mChy levels, with measurements taken every 15 minutes over a 14 hour time course. Strain AA639 performs the MiSTY cell differentiation program when stimulated with galactose, but 90% of the cell population remains in the ISC state when cultured in only glucose. Strain AA713 is an isolated population of Factory Cells derived from AA639. Strain AA090 is the background strain upon which the others were constructed. Bars show standard deviation among three independent trials.

### Growth Dynamics and Productivity

Because Switch #2 was designed to convert Factory Cells into leucine auxotrophs, we could compare culture growth rates and mChy synthesis in different defined media to determine the effectiveness of our strategy for decoupling the tasks of cell growth and product synthesis. We compared four different cultures: the BY4742 background strain grown in SDgal+glu, AA639 grown in SDglu (which is comprised of 90% ISCs), AA639 grown in SDgal+glu for 8 hours prior to start (comprised of approximately 50% ASCs, together with ISCs and Factory Cells), and AA713 grown in SDgal+glu (a Factory Cell isolate from AA639). Each culture was grown to exponential phase, then diluted to OD₆₀₀ 0.1 in a 96-well plate in leucine-deficient or leucine-supplemented media. We measured optical density and mChy fluorescence intensity every 15 minutes over a 14 hour growth period (Figure 8A-B).

In leucine-supplemented medium, all four strains exhibited comparable exponential growth rates and reached similar final densities, indicating that the genetic modifications in AA639 do not impair fitness when nutrients are available. ISC cultures accumulated a modest amount of mChy, owing to the basal level of Factory Cell differentiation in SDglu media. Compared to ISC-dominated cultures of AA639, galactose-stimulated ASC and Factory Cell-only cultures accumulated 2.5-fold higher levels of mChy, consistent with their being largely comprised of differentiated Factory Cells.

In leucine-deficient medium, growth profiles diverged sharply. Cultures containing the background strain and Factory Cells only failed to grow over the 14 hour time course, as expected for leucine auxotrophs. ISC cultures exhibited robust exponential growth, reflecting the activity of *LEU2*. ASC cultures, despite containing an increasing fraction of growth-inhibited Factory Cells, demonstrated sustained biomass accumulation that eventually slowed but still reached stationary phase by 14 hours. Since this culture could not have undergone more than 11 doublings since the initiation of ASC conversion, few of the ASCs would have expired from senescence over this time period, thus sustained growth is expected. Under these conditions, the accumulation of mChy in Factory Cell only and ISC cultures was quite low, owing to the total inhibition of cell growth in the former and the small percentage of Factory Cells in the latter. In contrast, the level of accumulated mChy in stationary phase galactose-stimulated ASC cultures was 4.5-fold higher than ISC-dominated cultures of AA639 and 9-fold higher than Factory Cell only cultures, demonstrating robust growth and mChy accumulation.

These results highlight a key feature of the MiSTY cell differentiation program: that biomass accumulation and mChy product synthesis can occur simultaneously, even under conditions that severely limit the growth and metabolic activity of Factory Cells. This validates the core design principle that division of labor can mitigate the conflicting needs of allocating resources towards cell growth or biosynthetic output.

## Discussion

In this work, we introduce MiSTY, a synthetic, lineage-based differentiation platform that brings microbial stem-cell architecture to *Saccharomyces cerevisiae*. It exploits the native asymmetry of budding yeast by using Ace2-mediated regulation of the p*CTS1* promoter to drive bud cell differentiation, which generates stable subpopulations and a genetically encoded division of labor. Sequential, irreversible recombinase-mediated rearrangements produce three defined cell states: (1) Inactive Stem Cells (ISCs) that proliferate, (2) Activated Stem Cells (ASCs) that self-renew while producing specialized progeny, and (3) Factory Cells (FCs) that terminally differentiate to execute engineered biosynthetic functions. This architecture restricts metabolic burden to FCs while ASCs sustain long-term population growth.

Quantitative lineage tracking and population assays show that ASCs reliably generate FC bud cells while maintaining their own proliferative capacity. Phenotypic assays and whole-genome sequencing confirm high differentiation fidelity and long-term genetic stability. By confining pathway activation to FCs, MiSTY supports sustained product synthesis without compromising ASC fitness, demonstrating reproductive and biosynthetic division of labor as a robust design principle for yeast biomanufacturing.

### Division of Labor Across Bacterial and Eukaryotic Systems

The successful implementation of MiSTY adds yeast to a growing set of microbial platforms that employ differentiation to resolve the growth–production trade-off. In bacteria, terminal differentiation systems such as the MiST platform in *E. coli* have shown that stem-like progenitor populations can generate short-lived, metabolically burdened producers while protecting the replicative lineage from production stress (Williams & Murray, 2022),(Glass *et al*, 2024),(Mushnikov *et al*, 2025). Similarly, bacterial co-culture approaches that divide muli-step biosynthetic pathways between different strains demonstrate that distributing the metabolic load can relieve burden and enhance yields, especially for toxic or energetically costly pathways (Grandel *et al*, 2025) (Flores *et al*, 2020), (Ma *et al*, 2021).

MiSTY adapts and extends these design principles to *Saccharomyces cerevisiae*. Budding yeasts provide a robust natural asymmetry that reliably distinguishes mother and daughter identities (Mazanka *et al*, 2008),(Herrero *et al*, 2020), offering a built-in mechanism for programming cell differentiation without reliance on artificially polarized systems. This advantage is complemented by yeast’s compatibility with complex eukaryotic post-translational modifications, compartmentalized metabolism, and widespread industrial use (Parapouli *et al*, 2020),(Kim *et al*, 2015), (De Wachter *et al*, 2021).

In synthetic yeast co-culture systems, distributing metabolic tasks across distinct subpopulations can stabilize growth and boost the production of complex molecules, particularly when strains are engineered to cross-feed essential metabolites (Peng *et al*, 2024),(Aulakh *et al*, 2023),(Roell *et al*, 2019). MiSTY captures this central benefit of co-culture systems while avoiding many of their operational challenges. By genetically encoding differentiation within a single isogenic population, MiSTY achieves an intrinsic division of labor in which growth and production are cleanly separated between subpopulations. Once activated, the system autonomously establishes and maintains its internal structure without the need for continuous external regulation, finely tuned inoculation ratios, or optimization of metabolite exchange. Self-organization allows MiSTY to retain the advantages of co-cultures while maintaining the simplicity of a single-strain platform.

### Limitations

Despite its strengths, MiSTY also carries several limitations. First, Factory Cell differentiation in MiSTY is not under positive selective pressure. During product synthesis, FCs may suffer metabolic burden and consequently become under-populated due to diminished proliferative capacity or even cell death, particularly when there are no selective forces that increase the number of Factory Cells (Rugbjerg *et al*, 2018). In contrast, cross-feeding consortia are designed to generate ecological dependencies in which each strain benefits from maintaining the community population structure, thereby providing selective reinforcement of division of labor (Peng *et al*, 2024), (Grandel *et al*, 2025).

Second, FCs in MiSTY shoulder the entire biosynthetic pathway load. While this simplifies genetic design, it contrasts with pathway-partitioned consortia, where distributing intermediates across strains can reduce metabolic stress, balance precursor fluxes, and enhance overall productivity (Ganesan *et al*, 2017). By placing all pathway steps into a single non-growing cell type, MiSTY may limit maximal production capacity for pathways that are highly burdensome or toxic.

Third, ASC identity is directed to mother cells, which divide approximately 25 times before senescing (Fehrmann *et al*, 2013), (Kaeberlein *et al*, 2005). As observed in metazoa, stem cell aging could be a barrier to long-term maintenance of the system (Oh *et al*, 2014). Hypothetically, an alternative configuration of the MiSTY design principle could employ a reversal of the polarity of asymmetric cell division, such that bud cell progeny are instructed to retain the ASC fate.

Although the MiSTY genetic system described in this work offers a stable single-strain alternative to complex synthetic consortia, subsequent versions of this technology may benefit from the incorporation of additional features. These may include selective incentives for Factory Cell proliferation and mechanisms that allow sub-specialization among differentiating bud cell progeny. The recombinase-mediated mechanisms that are built into the MiSTY cell differentiation pathway enable a broad range of opportunities for downstream genetic engineering.

## Materials and Methods

### Strains and Plasmids

All yeast strains were derived from BY4742 (MATα, his3Δ1, *Leu2*Δ0, lys2Δ0, ura3Δ0). Plasmid construction employed Gibson assembly using NEBuilder HiFi DNA Assembly Master Mix (New England Biolabs). Genomic integration vectors pYPRC15 and pYORW22 were constructed by amplifying 500 bp homology regions flanking target loci with appropriate selective markers (Ura3 or His3) and assembling into pMCS2 backbone. Complete strain genotypes and plasmid sequences are provided in Supplementary Tables S1-S2.

### Gene Construction and Genomic Integration

The pCTS1-Sic1deg-NES-GFP-Cre-NLS cassette was assembled by PCR amplification of pCTS1 (700 bp) and Sic1(amino acids 1-160) from BY4742 genomic DNA, GFP-Cre from AAV-Cre-GFP plasmid, and synthetic NLS/NES sequences. SV40 nuclear localization signal (NLS) PKKKRKV and nuclear export signal (NES) from a protein kinase inhibitor, LALKLAGLDI, followed by Gibson assembly into pYPRC15. The inverted circuit variant incorporated attB/attP sites flanking the inverted segment (pCTS1 position -335 to Cre codon 229) by sequential PCR and assembly. The pGAL1-Bxb1 construct was assembled by amplifying pGAL1 from BY4749 genomic DNA and Bxb1 coding sequence, then cloning into pYORW22. The loxP-flanked LEU2-mCherry reporter was synthesized (IDT gBlocks) and assembled into the

### HO-POLY-HO integration vector

For genomic integration, plasmids were linearized with NotI (pYPRC15, pYORW22) or EcoRV (HO-POLY-HO) and transformed into yeast using the lithium acetate method. Transformants were selected on appropriate dropout media and integration confirmed by colony PCR and sequencing. Integrations were directed to defined genomic loci (YPRC on chromosome XVI and HO on chromosome IV) at sites known to support expression of heterologous genes.

### Culture Media and Growth Conditions

Unless noted otherwise, overnight cultures were grown at 30°C in synthetic dropout medium (Sigma-Aldrich) with 0.4% glucose lacking leucine (Leu⁻). For ASC inductions, cultures at OD₆₀₀ ∼0.6 were pelleted, resuspended in fresh Leu⁻ medium containing 0.4% glucose and 2% galactose, and incubated at 30°C with shaking. Samples were collected at indicated timepoints for analysis.

For agar plating assays, cultures were diluted to OD₆₀₀ 0.1 and serially diluted over four rounds of 1:10 dilution. 5 µL was spotted onto leucine-deficient or leucine-supplemented plates. For growth in 96-well plates, cultures were diluted to OD₆₀₀ 0.1, placed in 96-well plates, and measured every 15 minutes over 14 hours at 30°C in a BioTek Synergy H4 plate reader.

### Microscopy

Time-lapse fluorescence microscopy was performed on an Axio Imager.Z2 microscope with Zen imaging software. GFP was detected using Filter set 38 HE (excitation BP 470/40, emission BP 525/50); mCherry using Filter set 63 HE (excitation BP 572/25, emission BP 629/62). For time-lapses, cells were mounted on 0.8% agarose pads containing appropriate medium and imaged every 15 minutes for 8 hours. Exposure times: 10 ms phase contrast, 300 ms GFP, 500 ms mCherry. ImageJ was used for image analysis and fluorescence intensity quantification following background subtraction.

### Flow Cytometry

Flow cytometry was performed on a BD FACS machine. mCherry fluorescence was measured in both fixed and live cells using appropriate filter sets. For sorting experiments, cells were separated based on mCherry intensity with >98% post-sort purity confirmed by reanalysis. Data were analyzed using Floreada.io software. For time-course experiments, cultures were induced as described and samples collected at 0, 4, 8, 12, 24, 48, and 72 hours for immediate analysis.

### Whole-Genome Sequencing

High molecular weight genomic DNA was extracted using the NEB Monarch Spin gDNA Extraction Kit. DNA quality was assessed using Agilent Genomic DNA ScreenTape. After native barcoding, gDNA samples were divided across two PromethION 2 flow cells for DNA sequencing (Oxford Nanopore), providing minimum and average total number of reads of 39,122 and 86,858 across datasets, respectively, and genomic coverage of 97X and 240X, respectively. Samtools was used to map sequencing reads against contigs derived from whole-genome reconstructions of sequence data from the T0 and T72 time points, which were confirmed to have both switch elements of the MiSTY genetic system in the initial and final states, respectively. Integrated Genomics Viewer was used to observe sequence alignments at the MiSTY switch element loci. For each dataset, reads that spanned the inversion or deletion target loci, included at least 300 bp of sequence within the inversion or deletion target loci, or read more than 300 bp beyond the deletion site scar, were manually identified and counted. To give datasets equal weight, raw counts were expressed as a fraction of the total number of reads per dataset and normalized with respect to the dataset with the highest total number of reads.

## Author Contributions

## Acknowledgments

We thank Nic Blouin of the Wyoming INBRE Bioinformatics Core for assistance with genome sequencing and analysis. Mark Gomelsky, Nik Mushnoikov, and Steven Poyer provided insightful suggestions that advanced the work. GB was supported by a research grant from the National Science Foundation (2225849), and both GB and JG were supported by Small Business Innovation and Research grants from the National Institutes of Health (1R41GM137710) and the National Science Foundation (2222602), by the Wyoming Business Council, and by an Institutional Development Award (IDeA) from the National Institutes of Health under Grant # 2P20GM103432.

