## Supplementary Figures and Tables for "Programmable cell differentiation in budding yeast uncouples reproductive and metabolic tasks"

### Supplementary Figure 1

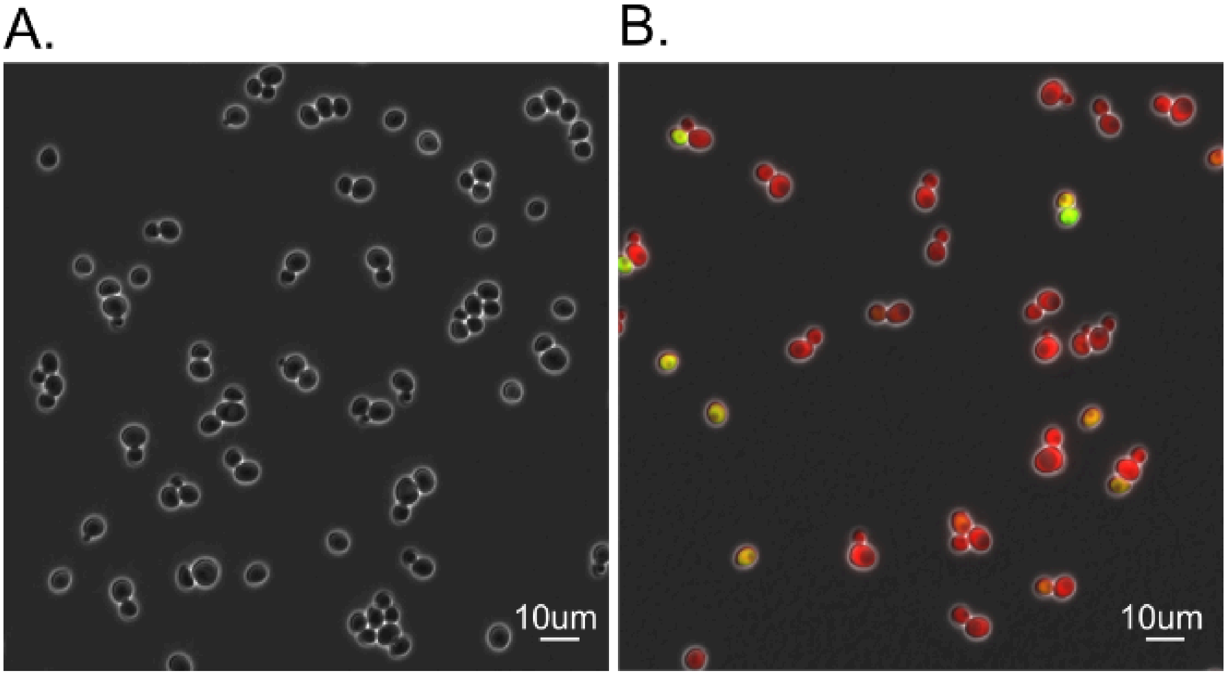


**Supplementary Figure 1 | Functionality of Cre Recombinase Reporter Gene**

A) Fluorescence microscope image of Strain AA320 (Cre recombinase reporter integrated into the BY4742 genetic background). B) Fluorescence microscope image of Strain AA321 (p*CST1*-Cre and the Cre recombinase reporter integrated into BY4742). In both images, signals from GFP (green) and mChy (red) channels are overlayed on the phase contrast image (gray).

#### Supplementary Figure 2

A.

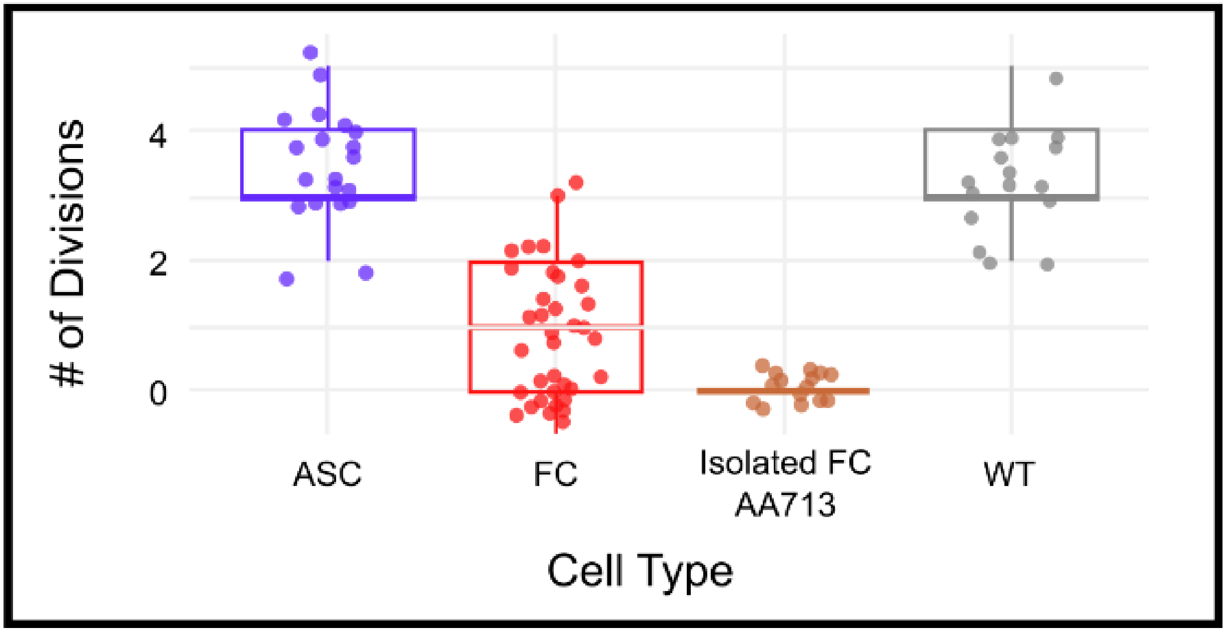


**Supplementary Figure 2 | Capacity For Cell Division in Leucine-Deficient Media**

Quantification of the number of cell divisions completed by individual cells over 8-hours of incubation during microscope time-lapse experiments. ASC: Strain AA639 was cultured on SDglu+gal -Leu media, and the experiment follows the divisions of non-fluorescent activated stem cells. FC: Strain AA639 was cultured on SDglu+gal -Leu media, and the experiment follows the divisions of newborn Factory Cells. Strain AA713: A Factory Cell isolate from Strain AA639 was cultured on SDglu+gal – Leu media. WT: BY4742 genetic background.

#### Supplementary Figure 3

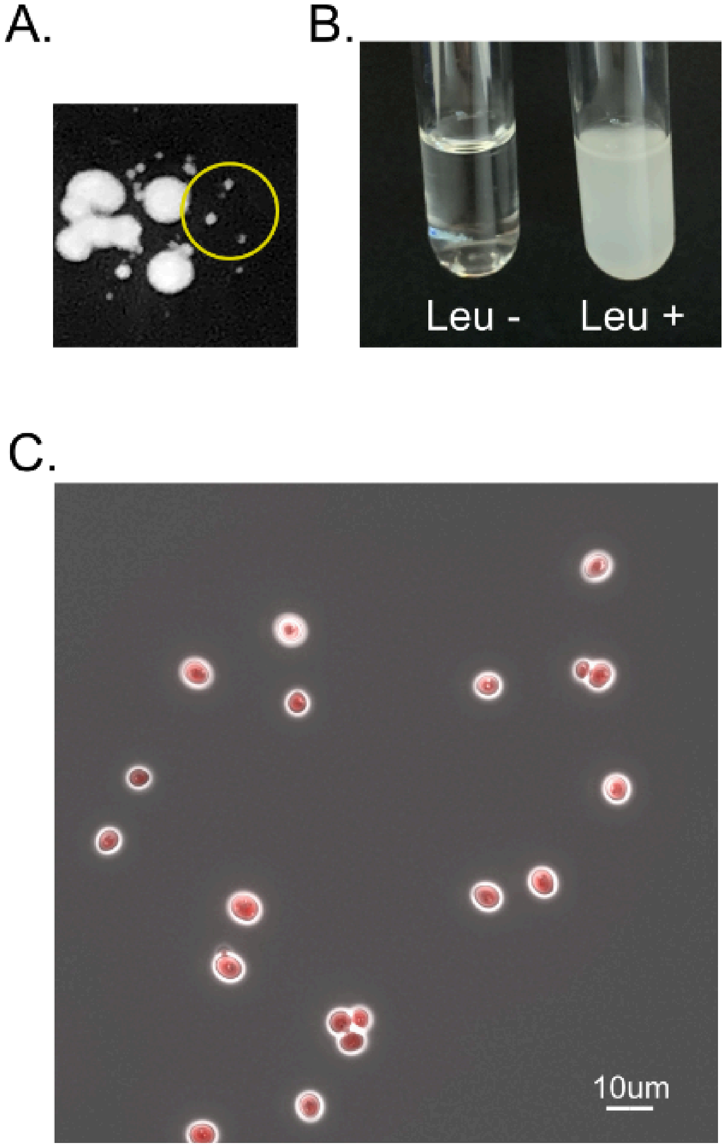


**Supplementary Figure 3 | Micro-colony Analysis**

A) A close-up image of a spot-plated area from the agar plate in Figure 6A, revealing micro-colonies (small white dots, shown in yellow circle). B) Overnight growth of a micro-colony inoculated into leucine-deficient (Leu-) or leucine-supplemented liquid media (Leu+). C) Fluorescence microscopy of a micro-colony grown in Leu+ media.

**Supplementary Video V1**: Time-lapse video microscopy of Strain AA311. This video shows the cell cycle-dependent localization of the Cre fusion protein in Strain AA311, which contains the p*CTS1*-Cre gene in a normal BY4742 genetic background. GFP signal (green) is overlayed on the phase contrast image (gray). The data shown in Figure 2B was obtained from this video.

**Supplementary Video V2**: Time-lapse video microscopy of Strain AA639. This video shows the cell cycle-dependent localization of the Cre fusion protein and subsequent accumulation of mChy in Strain AA639, which contains the complete MiSTY genetic system. Signals from the GFP (green) and mChy (red) channels are overlayed on the phase contrast image (gray). The data shown in Figure 4A was obtained from this video.

**Supplemental Table 1 | Strains in this Work**

| Strain | Species | Description | Source | Marker |
|---|---|---|---|---|
| A090 | *S. cerevisiae* | BY4742 (MATα his3Δ1 Leu2Δ0 lys2Δ0 ura3Δ0) | Eunsook Park | |
| AA320 | *S. cerevisiae* | Cre reporter - BY4742 (MATα his3Δ1 lys2Δ0 ura3Δ0), HO :: pTet - Leu2 - mCherry | This Study | Leu |
| AA104 | *S. cerevisiae* | BY4742 with plasmid pCM109-SV40NLS-mCherry PopZ | This Study | Ura |
| AA105 | *S. cerevisiae* | BY4742 with plasmid pCM190-mCherry-PopZ | This Study | Ura |
| AA311 | *S. cerevisiae* | BY4742, YPRC :: pCST1-Sic1(1-160)-NES-GFP-Cre-NLS | This Study | Ura |
| AA321 | *S. cerevisiae* | Cre reporter, YPRC :: pCST1-Sic1(1-160)-NES-GFP-Cre-NLS | This Study | Ura, Leu |
| AA489 | *S. cerevisiae* | BY4742, YPRC :: pCST1-Sic1(1-160)-NES-GFP-Cre (1-270)-GID1-NLS | This Study | Ura |
| AA490 | *S. cerevisiae* | Cre reporter, YPRC :: pCST1-Sic1(1-160)-NES-GFP-Cre (1-270)-GID1-NLS | This Study | Ura, Leu |
| AA503 | *S. cerevisiae* | BY4742, YORW :: pGal01-GAI-Cre (271-343)-NLS | This Study | His |
| AA505 | *S. cerevisiae* | Cre reporter, YPRC :: pCST1-Sic1(1-160)-NES-GFP-Cre (1-270)-GID1-NLS, YORW :: pGal01-GAI-Cre (271-343)-NLS | This Study | Ura, His, Leu |

| | | | | |
|---|---|---|---|---|
| AA532 | *S. cerevisiae* | BY4742, YORW :: pGal01-GFP | This Study | His |
| AA533 | *S. cerevisiae* | BY4742, YORW :: pGal01-(stronger RBS)-GFP | This Study | His |
| AA548 | *S. cerevisiae* | Cre reporter, YPRC :: pCST1-Sic1(1-160)-NES-GFP-Cre (1-270)-GID1-NLS, YORW :: pGal01-(stronger RBS)-GAI-Cre (271-343)-NLS | This Study | Ura, His, Leu |
| AA592 | *S. cerevisiae* | Cre reporter, YPRC :: pCST1-Sic1(1-160)-NES-GFP-Cre-NLS, Nucleotides (584-420) upstream of the transcriptional start were inverted with flanking Bxb1 *attP* and *attB* sites. | This Study | Ura, Leu |
| AA593 | *S. cerevisiae* | Cre reporter, YPRC :: pCST1-Sic1(1-160)-NES-GFP-Cre-NLS, Nucleotides (584-420) upstream of the transcriptional start were inverted with flanking Bxb1 *attP* and *attB* sites. YORW :: pGal01-(stronger RBS)-Bxb1 | This Study | Ura, His, Leu |
| AA610 | *S. cerevisiae* | BY4742, YPRC :: pCST1-Sic1(1-160)-NES-GFP-Cre-NLS, Bxb1 *attP* inserted after AA229 in Cre | This Study | Ura |
| AA611 | *S. cerevisiae* | BY4742, YPRC :: pCST1-Sic1(1-160)-NES-GFP-Cre-NLS, Bxb1 *attP* inserted after AA251 in Cre | This Study | Ura |
| AA636 | *S. cerevisiae* | BY4742, YPRC :: pCST1-Sic1(1-160)-NES-GFP-Cre-NLS, Inverted fragment 335bp | This Study | Ura |

| | | | | |
|---|---|---|---|---|
| | | upstream of transcriptional start through aa229 in Cre. Inverted region flanked by Bxb1 *attB* and *attP* sites. | | |
| AA639 (MiSTY) | *S. cerevisiae* | Cre reporter, YPRC :: pCST1-Sic1(1-160)-NES-GFP-Cre-NLS, Inverted fragment 335bp upstream of transcriptional start through aa229 in Cre. Inverted region flanked by Bxb1 *attB* and *attP* sites, YORW :: pGal01-(stronger RBS)-Bxb1 | This Study | Ura, His, Leu |
| AA713 | *S. cerevisiae* | Cre reporter, YPRC :: pCST1-Sic1(1-160)-NES-GFP-Cre-NLS, Inverted fragment 335bp upstream of transcriptional start through aa229 in Cre. Inverted region flanked by Bxb1 *attB* and *attP* sites, YORW :: pGal01-(stronger RBS)-Bxb1. (FC control) | This Study | Ura, His, Leu |
| AA715 | *S. cerevisiae* | Cre reporter, YPRC :: pCST1-Sic1(1-160)-NES-GFP-Cre-NLS, Inverted fragment 335bp upstream of transcriptional start through aa229 in Cre. Inverted region flanked by Bxb1 *attB* and *attP* sites | This Study | Ura, Leu |

**Supplemental Table 2 | Plasmids in this Work**

| Plasmid Name | Source | Essential Genes and Regulatory Elements | Selective Marker |
|---|---|---|---|
| pMCS2 | Martin Thanbichler MTLS collection | pMCS2 | Kan |
| pYPRC15 | This Study | YPRC homology (500bp)-MCS-Ura3-YPRC homology (500bp) | Kan |
| pYORW22 | This Study | YORW homology (500bp)-MCS-His3-YORW homology (500bp) | Kan |
| HO-poly-HO | HO-Poly-HO was a gift from David Stillman (Addgene plasmid # 51661 ; http://n2t.net/addgene:51661 ; RRID:Addgene_51661) | HO homology (725bp)-MCS-HO homology (387bp) | Amp |
| AAV-Cre-GFP | AAV-Cre-GFP was a gift from Eric Nestler (Addgene plasmid # 68544 ; http://n2t.net/addgene:68544 ; RRID:Addgene_68544) | Cre fused with EGFP | Amp |
| pYTK076 | Eunsook Park | His3 marker | |
| pCM190-mCherry-PopZ | This Study | pCM 190 + mCherry-PopZ | Amp |

| | | | |
|---|---|---|---|
| pCM190-SV40NLS-mCherry-PopZ | This Study | pCM 190 + SV40 NLS_mCherry-PopZ | Amp |
| pBad-CSGC | This Study | pBad_CST1 promoter-Sic1(1-160)-NES-GFP-Cre-NLS,Tsynth25 | Amp |
| pUCIDT-YPRC_MCS_Ura3 | This Study | YPRC integration_MCS_Ura3 with pUCIDT-Amp+ | Amp |
| pMCS2-CSGC | This Study | pMCS2-CST1 promoter-Sic1(1-160)-NES-GFP-Cre-NLS,Tsynth25 | Kan |
| pUCIDT-YPRC-CSGC-Ura3 | This Study | YPRC homology (500bp)-CST1 promoter-Sic1(1-160)-NES-GFP-Cre-NLS,Tsynth25-Ura3-YPRC homology (500bp) | Amp |
| HO-XLXM-HO | This Study | HO homology (725bp)-loxP-Leu2-ADH1 terminator-loxP-mCherry-HO homology (387bp) | Amp |
| pBW2426 | pBW2426_pCAG-PV1-iCre-N270-L1-GID1-NLS-BGHpA was a gift from Wilson Wong (Addgene plasmid # 108730 ; http://n2t.net/addgene:108730 ; RRID:Addgene_108730) | pCAG-PV1-iCre-N270-L1-GID1-NLS-BGHpA | Amp |
| pBW2436 | pBW2436_pCAG-PV1-GAI-L1-Cre-271C-NL | pCAG-PV1-GAI-L1-Cre-271C-NLS-BGHpA | Amp |

| | | | |
|---|---|---|---|
| | S-BGHpA was a gift from Wilson Wong (Addgene plasmid # 108730 ; http://n2t.net/addgene:108730 ; RRID:Addgene_108730) | | |
| pYPRC15_CSGCsplitN270 | This Study | YPRC homology (500bp)-pCST1-Sic1(1-160)-NES-GFP-Cre (1-270)-L1-GID1-NLS-Ura3-YPRC homology (500bp) | Kan |
| pYORW22_pGal-GIA-Crec271 | This Study | YORW homology (500bp)-pGal01-GAI-Cre (271-343)-NLS-His3-YORW homology (500bp) | Kan |
| pYORW22_GFP | This Study | YORW homology (500bp)-pGal01-GFP-His3-YORW homology (500bp) | Kan |
| pYORW22_RBS01-GFP | This Study | YORW homology (500bp)-pGal01(stronger RBS)-GFP-His3-YORW homology (500bp) | Kan |
| pYORW22_Gal1_RBS_C271 | This Study | YORW homology (500bp)-pGal01(stronger RBS)-GAI-Cre (271-343)-NLS-His3-YORW homology (500bp) | Kan |
| pYORW22-pGal1-BXB1 | This Study | YORW homology (500bp)-pGal01(stronger RBS)-Bxb1-His3-YORW homology (500bp) | Kan |

| | | | |
|---|---|---|---|
| pYPRC15-Cre-AA229 *attP* | This Study | YPRC homology (500bp)-pCST1-Sic1(1-160)-NES-GFP-Cre-NLS, Bxb1 *attP* inserted after AA229 in Cre-Ura3-YPRC homology (500bp) | Kan |
| pYPRC15-Cre-AA251 *attP* | This Study | YPRC homology (500bp)-pCST1-Sic1(1-160)-NES-GFP-Cre-NLS, Bxb1 *attP* inserted after AA251 in Cre-Ura3-YPRC homology (500bp) | Kan |
| pYPRC15-MiSTY | This Study | YPRC homology (500bp)-pCST1-Sic1(1-160)-NES-GFP-Cre-NLS, Inverted fragment 335bp upstream of transcriptional start through aa229 in Cre. Inverted region flanked by Bxb1 *attB* and *attP* sites-Ura3-YPRC homology (500bp) | Kan |
